# PlantNexus: A Gene Co-expression Network Database and Visualization Tool for Barley and Sorghum

**DOI:** 10.1101/2021.04.23.441196

**Authors:** Yadi Zhou, Abhijit Sukul, John W. Mishler-Elmore, Ahmed Faik, Michael A. Held

## Abstract

Global gene co-expression networks (GCNs) are powerful tools for functional genomics whereby putative functions and regulatory mechanisms can be inferred by gene co-expression. With the recent accumulation of RNA-seq data sets, the construction of RNA-seq-based GCNs has now become possible. Cereal crops, such as *Hordeum vulgare* (barley) and *Sorghum bicolor* (sorghum), are among the most important plants to humanity and contribute significantly to our food supply. However, co-expression network tools for these plants are outdated or lacking. In this study, we constructed global GCNs for barley and sorghum using 500 and 774 RNA-seq data sets, respectively. In addition, we curated the meta-information of these RNA-seq data sets and categorized them into four main tissue types, leaf, root, shoot, and flower/seed, and built tissue-specific GCNs. To enable GCN searching and visualization, we implemented a website and database named PlantNexus, offering an immersive environment for the exploration and visualization of gene expressions and co-expressions of barley and sorghum at the global and tissue-specific levels. PlantNexus is freely available at https://plantnexus.ohio.edu/.

## Introduction

Next-generation RNA sequencing (RNA-seq) technology has become the preferred way of generating gene expression profiles in a high-throughput manner. RNA-seq data sets encode important information of cellular processes. These transcriptomic data sets can be decoded for many purposes with appropriate techniques. Large-scale RNA-seq data can be used to study gene functions and regulatory mechanisms (1, 2) through gene co-expression networks (GCNs). GCNs allow simultaneous identification and classification of many genes with similar expression characteristics (2). For example, GCNs have been used to understand the genetic basis of plant natural products (3), nitrogen metabolism for plant growth (4), and cell wall development (5).

GCNs have been constructed for several plants, such as rice (6, 7), maize (8), wheat (9), soybean (10). Several plant co-expression databases have been published, for example, ATTED-II (11), RiceFREND (7), PLANEX (12), PlaNet (13), and BarleyNet (14) for the exploration of gene co-expressions. However, GCN tools are limited by outdated datasets or are not currently available for some plant species, for example, *Sorghum bicolor* (sorghum). Barley and sorghum are among the most important cereal crops to the humanity. They are used for human diet, livestock feed, and alcoholic beverage production, etc. It is estimated that a 70% increase in agricultural output will be required to feed the world’s population by 2050 (15). A better understanding of gene functions and regulatory mechanisms is essential for developing strategies to increase grain yield and nutrient qualities. Original barley GCNs were constructed using microarray-based data (16), were local GCNs constructed from certain conditions or treatments (17, 18), or were constructed using relatively few RNA-seq data (14, 19).

In this study, we used all available high-quality RNA-seq-based transcriptomic data sets across various tissue types, developmental stage, and treatment conditions to construct global GCNs for barley and sorghum. These GCNs are freely accessible through a web interface with interactive visualization tools hosted at PlantNexus at https://plantnexus.ohio.edu/ (**Fig. 1A**).

**Figure 1.**
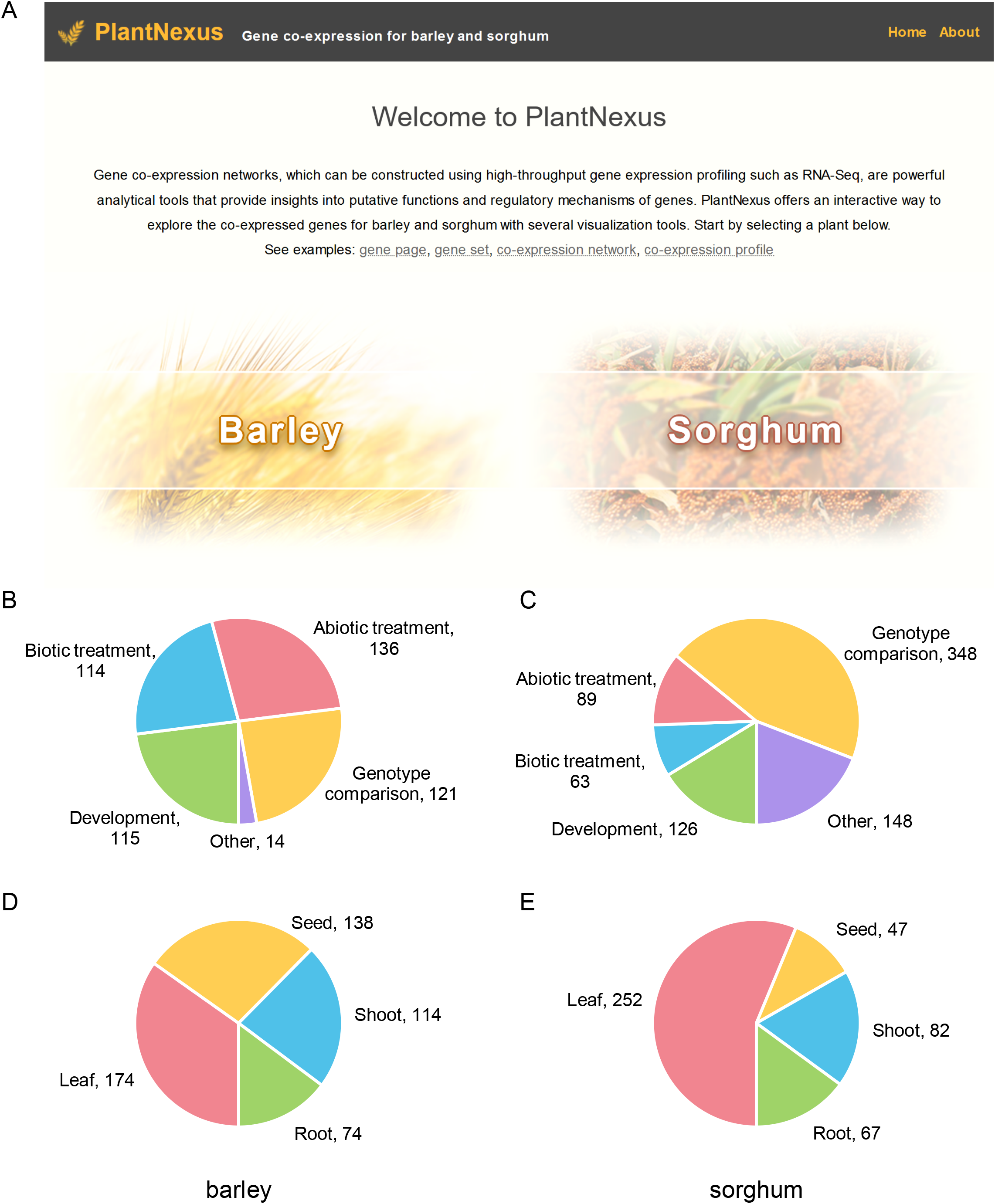
PlantNexus is a new web service for querying and visualizing gene expression and co-expression for barley and sorghum. (**A**) The home page of PlantNexus. (**B**) Study type distribution for the barley RNA-seq data sets in PlantNexus. (**C**) Study type distribution for the sorghum RNA-seq data sets in PlantNexus. (**D**) Tissue type distribution for the barley RNA-seq data sets in PlantNexus. (**E**) Tissue type distribution for the sorghum RNA-seq data sets in PlantNexus. Number shows the RNA-seq data sets for the corresponding category. Data sets with known study types or tissue types are included in the distribution plots.

## Materials and Methods

### RNA-seq data collection

We queried the NCBI SRA database for RNA-seq data sets for barley (on January 2018) and sorghum (on January 2019) using the term “(hordeum_vulgare[Organism] OR sorghum_bicolor[Organism]) AND (biomol_transcript[properties] OR study_type_transcriptome_analysis[properties] OR strategy_rna_seq[properties] OR strategy_FL_cDNA[properties])”. We identified transcriptomic data sets that utilized an RNA-seq strategy. We also manually extracted meta-information including tissue types, developmental stages, and treatments, etc., based on the NCBI SRA database and the original publications of these data sets. A total of 735 barley RNA-seq data sets and 1225 sorghum RNA-seq data sets were downloaded.

### RNA-seq data processing pipeline

SRA files were converted to FASTQ format using the fastq-dump command from the SRA toolkit v2.8.2. The quality of raw RNA-seq data sets was examined using FastQC v0.11.5 (20). Low quality samples, such as those with low average quality scores and overrepresented sequences were excluded. All data sets were trimmed by Trimmomatic v0.36 (21) using default parameters “LEADING:3 TRAILING:3 SLIDINGWINDOW:4:15 MINLEN:36”. All trimmed data sets were mapped to barley genome v36 and sorghum v41 from Ensembl Plant (22) using STAR v2.5.3 (23) with default parameters “--quantMode GeneCounts --outSAMtype None --outSAMmode None -- readFilesCommand zcat”. Genome indexes for mapping with STAR were generated using parameters “--sjdbOverhang 100 --genomeChrBinNbits 14 -- genomeSAindexNbases 12 --genomeSAsparseD 3”. Gene-level expression read counts were directly generated by STAR.

RNA-seq data sets that met the following criteria were used for the construction of barley GCNs (24): (1) that >70% of the reads survived after trimming; (2) that >70% of the reads mapped to the barley genome uniquely; (3) that they contain at least 10 million uniquely mapped reads. A total of 500 barley data sets and 774 sorghum data sets met these criteria. Raw gene count was normalized using transcripts per kilobase million (TPM) and log_2_ transformed.

### Co-expression network construction

Next, we constructed a global GCN using these selected RNA-seq data sets. Tissue-specific GCNs were also constructed using RNA-seq data sets for specific tissue types for barley and sorghum. To ensure the genes being evaluated for co-expression had adequate expression levels, we filtered the genes and kept those that had a count per million (CPM) ≥ 1 in at least 20 data sets, which resulted in 23728 barley genes 26638 sorghum genes. The expression data for all individual genes, however, are still available in PlantNexus.

Gene co-expression was computed using the widely used method, Pearson correlation coefficient (PCC). We computed a PCC score for each gene pair.

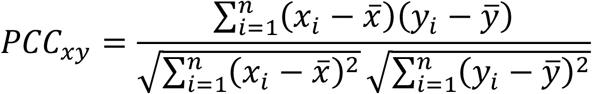

The process was repeated 1000 times, each time randomly using 80% of the data sets. The PCC scores of the 1000 repeats were averaged to get a final correlation for each gene pair. The PCC scores were further transformed by mutual rank (MR) (25):

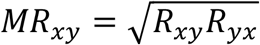

where *R*_*xy*_ is the rank of gene y in gene x’s *PCC*_*x*._ list, and *R*_*yx*_ is the rank of gene x in gene y’s *PCC*_.*y*_ list. The MR scores were used to rank the co-expressions of genes.

### Web service implementation

PlantNexus was implemented with Python (3.8) (https://www.python.org/) framework Django (3.1) (https://www.djangoproject.com/) for the server backend. For the database, SQLite (https://www.sqlite.org/) was used. The frontend was implemented with HTML, CSS, and JavaScript. The website was designed in a way that any data query to the database was transferred to the frontend by AJAX asynchronously in JSON format. This provides a smooth experience with highly interactive visualization tools and enables users to access all our data through user programs so that PlantNexus can be integrated into their pipelines. Network visualization was implemented using Cytoscape.js (26). PlantNexus is hosted by the Ohio University.

## Results and discussion

### Overall design of PlantNexus

PlantNexus is focused on the global expression and co-expression patterns of genes for barley and sorghum. To achieve this, we queried the NCBI SRA database and retrieve RNA-seq based transcriptomic profiles for barley and sorghum. After processing a large number of relevant RNA-seq data sets (see **Materials and methods**), we identified 500 data sets for barley and 774 data sets for sorghum. We manually curated meta information for these data sets, and categorized them into four major tissue types, namely leaf, seed, shoot, and root. We examined the distributions in terms of study types (**Fig. 1B-C**) and tissue types (**Fig. 1D-E**). The major study types were biotic treatment, abiotic treatment, developmental, and genotype comparison. For barley, these study types had similar numbers of data sets. For sorghum, genotype comparison accounted for 45% of the RNA-seq data sets we retrieved and processed. In terms of tissue types, leaf was the most popular tissue type for both barley and sorghum. Yet, all tissue types had an adequate number of data sets, with the minimum being seed for sorghum (47 data sets). This enabled us to simultaneously construct tissue-specific GCNs while building global GCNs using all data sets.

To build GCNs, we computed a co-expression score for each gene pair using PCC. The computation was repeated 1000 time, each time using 80% randomly selected data sets. The results were averaged. After we obtained an average PCC score for each gene pair, we transformed the scores using MR. MR considers the rank of two genes in each other’s list ranked by PCCs and is the geometric average of the two ranks. MR is the default co-expression ranking methods in many co-expression databases, such as ATTED-II (11) and RiceFREND (7). All results were integrated into a relational database, which includes the expression of each gene across the data sets, top co-expressed genes for each gene, basic information such as gene ontology terms extracted from the genome annotation, and percent of data sets that express a certain gene, etc. The expression and co-expression results are available in the global level (using all data sets) and tissue level (using corresponding data sets).

Compared to other platforms, PlantNexus has several strengths: (1) a large amount of RNA-seq data sets were processed with the same pipeline in a consistent manner; (2) in addition to the global GCNs, PlantNexus offers both GCN and tissue level co-expression results; and (3) PlantNexus has a highly centralized web interface where all data can be easily navigated, and interactive visualizations for the GCNs and expression levels.

### Web interface and visualizations

To serve the data, we implemented a web service named PlantNexus. The entry point to the data are “single gene” and “multiple gene” search (**Fig. 2A-B**), which is similar to other plant gene co-expression databases. However, one of the key strengths is that the web interface is highly centralized, meaning that users will be able to easily navigate to and keep track of the data and visualizations. This eliminates the need to switch between browser tabs. On the left side of the web page, users can find search boxes (**Fig. 2B**) and result list (**Fig. 2C**). The right-side area is organized by four tabs (**Fig. 2D**), “Getting Started”, “Data Table”, “Expression”, and “Network”. The “Data Table” shows the data table for a single gene or a gene set. “Expression” and “Network” contain visualization tools. When users click any internal links or buttons, the results will be loaded directly in the same web page and a new button will be added in the result list on the left. The users can switch between previous results and newly opened results using the buttons in the result list. The buttons are organized in tree forms such that the browsing order is preserved (**Fig. 2C**). Data tables and visualization tools are explained using two examples in the following sections.

**Figure 2.**
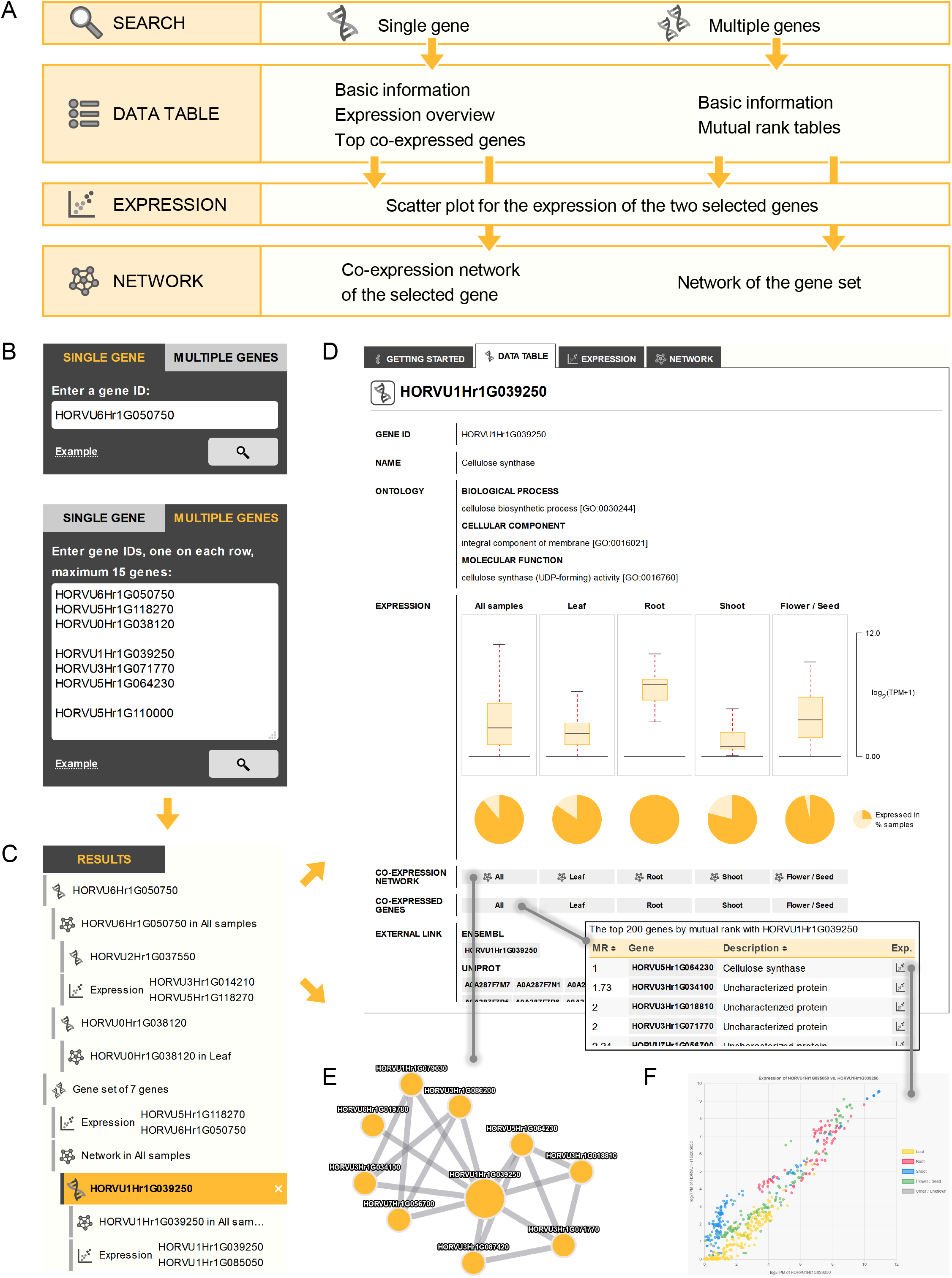
Overall design and web interface of PlantNexus. (**A**) Information architecture of PlantNexus. (**B**) Two search modes are available, “single gene” and “multiple genes”. (**C**) The search results and data loaded through user interactivities are organized in button trees. By clicking these buttons, users can switch the display of different results in the same web page. (**D**) An example of the data table for a single gene. For each gene, the average expression (box plots) and percent of data sets that express it (pie plots) are visualized. Co-expressed genes are sorted by the mutual rank (MR) in ascending order. By clicking the corresponding buttons from this page, users can view the co-expression network (**E**) and expression plot (**F**). The network viewer allows users to export the networks, change node and edge color, load selected node (gene data table) and edge (expression plot), annotate the node, and filter the network by various MR cutoffs. The expression plot allows users to filter the plot by tissue types.

### Example-1: single gene search

A data table of a single gene, such as the one shown in **Fig. 2D**, is displayed when users click any gene ID in PlantNexus. Here, the barley gene HORVU1Hr1G039250 is used as an example. This gene is also known as *Hordeum vulgare* cellulose synthase gene A4 (*HvCesA4*), which participates in cellulose synthesis in the secondary cell wall of barley (27). A query of HORVU1Hr1G039250 returns basic information including its name, gene ontology terms, and external links to Ensembl (22) and Uniprot (28). It also has box plots that represent the expression level of HORVU1Hr1G039250 in all samples and across individual tissue types and pie plots that represent the percentage of data sets that show *HvCesA4* expression. From the latter, we see that *HvCesA4* is expressed in all root data sets and in a high percentage of other tissue type data sets (>78.9%) (**Fig. 2D**, pie plots). The box plots we see that the overall expression of *HvCesA4* in root data sets (medium log_2_(TPM+1) = 7.01) is also higher than other tissues. These results are consistent with previous qPCR observations (27). By clicking the “All” button in the “Co-expressed genes” section, users can view the top co-expressed genes with *HvCesA4* among all datasets sorted by MR. A data table can be exported (.csv) by scrolling to the bottom of the table. As expected, among the top five genes co-expressed with *HvCESA4* are HORVU3Hr1G071770 (*HvCesA7*, MR=2) HORVU5Hr1G064230 (*HvCesA8*, MR=1), which are thought to participate in protein:protein interactions to form the barley secondary cell wall cellulose synthase complex (27). The same analysis can be performed using individual tissue type datasets by clicking either “leaf”, “root”, “shoot”, or “flower/seed”.

Clicking the buttons next to the “Co-expression network” section will open the GCNs for *HvCesA4* in the “Network” tab (**Fig. 2E**). The network viewer provides basic functions such as exporting as a network file (which can be loaded into software such as Cytoscape) or an image file. Layout, changing node / edge color, and changing node can all be modified. Users can load a specific node, which opens a new data table for the gene, or an edge, which opens the expression plot for the two connecting genes. Importantly, the network can be filtered by MR cutoff (e.g., MR≤5, MR≤10, MR≤20,…). This function allows the user to filter the network to show only the most co-expressed gene pairs. A table below the network lists all the co-expressions in the network with MR values. In addition to the network viewer, users can also visualize the expression of a selected co-expressed gene and *HvCesA4* (**Fig. 2F**) by clicking the plot button in the co-expression table or by loading an edge in the GCNs of *HvCesA4*. The tissue types are color-coded in the expression plot and their visibility can be controlled separately.

### Example-2: gene set search

To demonstrate a multiple gene search, the following genes were queried as an example: HORVU6Hr1G050750 (*HvCesA1*), HORVU5Hr1G118270 (*HvCesA2*), HORVU0Hr1G038120 (*HvCesA6*), HORVU1Hr1G039250 (*HvCesA4*), HORVU3Hr1G071770 (*HvCesA7*), HORVU5Hr1G064230 (*HvCesA8*), and HORVU5Hr1G110000 (*HvCesA3*). The data table tab for a set of genes contains a link to each gene and its description. An arrayed MR table for all data sets and each tissue type, can be opened by clicking the corresponding button (**Fig. 3A**). In the MR table, the length of the bar under each MR value is inversely proportional to the value. Clicking an individual cell opens the associated expression plot (**Fig. 3B**). The bottom button (Plot the network of these genes) allows one to plot the network for the gene set (**Fig. 3C**). As expected, the selected genes form two distinct networks. Cellulose synthesis in vascular plants is catalyzed by cellulose synthase complex in a hexameric rosette structure consists of multiple *CesA* genes (29). In barley, *HvCesA1, HvCesA2*, and *HvCesA6* are responsible for cellulose synthesis in primary cell wall, while *HvCesA4, HvCesA7*, and *HvCesA8* are responsible for secondary wall cellulose synthesis (27). The MR table shows high co-expression (low MR value) among the first group of *HvCesA1, HvCesA2*, and *HvCesA6* and the second group of *HvCesA4, HvCesA7*, and *HvCesA8*, yet there is no co-expression among these two groups (**Fig. 3A,C**). *HvCesA3* has a different expression pattern from these genes and is not a component of these two groups (27). The results indeed show that *HvCesA3* has a high MR (> 1000) with all other six genes (**Fig. 3A**) and is absent from the network (**Fig. 3C**). These results are consistent with previous publications.

**Figure 3.**
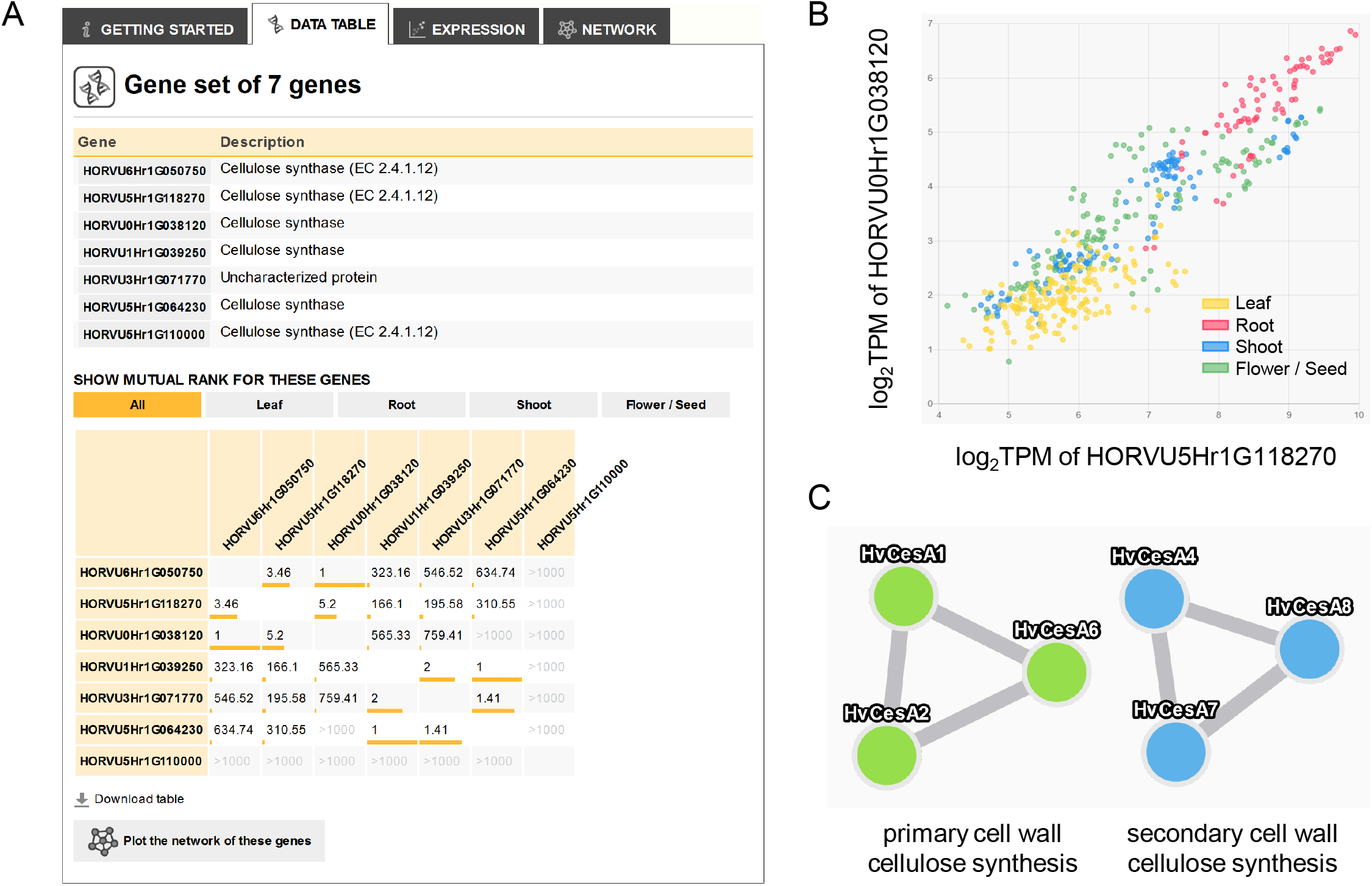
Example use of PlantNexus for gene set as input. (**A**) A mutual rank table is displayed when users query a set of genes. (**B**) Expression plot can be loaded directly by clicking the mutual rank in the table. (**C**) The co-expression of the genes in the gene set can be visualized in a network. In this example, cellulose synthesis genes in primary cell wall and secondary cell wall are disconnected among the two groups, but fully connected within groups.

## Conclusions

In this study, we present PlantNexus, a new web service and database for gene co-expression exploration for barley and sorghum, using a large number of high-quality RNA-seq data sets across many tissue types, developmental stages, and treatment conditions. PlantNexus will be updated annually or when a large number of new data sets are available and add new features based on user feedback. We believe PlantNexus will be a valuable resource for the plant research community focusing on barley and sorghum due to 1-the large amount of expression and co-expression data it hosts, 2-consistent RNA-seq processing pipelines used for all data sets, 3-manually curated rich meta information which enables tissue-level co-expression analyses, and 4-a highly centralized and interactive web interface.

## Declarations

### Availability of data and materials

All the SRA files are found in the NCBI SRA database. All the data in PlantNexus can be freely accessed without registration requirement at https://plantnexus.ohio.edu/.

### Authors’ contributions

YZ and MH conceived the study. MH and AF supervised the study. YZ constructed the database and developed the website. YZ, AS, and JE performed data gathering and processing. YZ and MH wrote the manuscript, and all authors critically revised the manuscript and gave final approval.

## Acknowledgements

We thank the Ohio Supercomputer Center for providing computing resources. We thank Ohio University OIT for hosting PlantNexus.

